# Genetic diversity and conservation insights for the baobab *Adansonia suarezensis* in northern Madagascar: a whole genome SNP analysis

**DOI:** 10.1101/2025.01.31.635890

**Authors:** Onja Hariveloniaina Morilline Razanamaro, Dario La Montagna, Fabio Attorre, Elisa Rondoni, Francesco Buttarazzi, Tahiana Andriaharimalala, Naoko Ishikawa, Yoshihisa Suyama, Laura Parducci

**Author notes:** Joint first authors (Onja Hariveloniaina Morilline Razanamaro, Dario La Montagna).

## Abstract

Madagascar is home to seven unique baobab species in the genus *Adansonia*, all threatened by climate change, habitat destruction, and deforestation. Previous studies have highlighted the vulnerability of baobabs, particularly *A. suarezensis*, without delving into the specific genetic structure of its populations. In this paper we examine the genetic structure, diversity, and connectivity of seven populations of *A. suarezensis*, using genome-wide Single Nucleotide Polymorphism (SNP) data. The results revealed significant genetic differentiation between inland and coastal populations, with the Mahory population forming a distinct genetic cluster characterized by high heterozygosity but low polymorphism, indicative of historical bottlenecks. In contrast, northern populations showed greater admixture and gene flow but higher inbreeding coefficients in coastal regions, such as Ambilo and Cap d’Ambre, suggesting localized inbreeding depression. Additionally, historical climatic shifts and potential anthropogenic dispersal are explored as contributing factors to the current genetic patterns. This study highlights the importance of understanding genetic dynamics for conservation, emphasizing habitat restoration and targeted management strategies to preserve the evolutionary potential of *A. suarezensis* and ensure its long-term survival.

## Introduction

Baobabs, iconic trees belonging to the genus *Adansonia* endemic to Madagascar, are under severe threat from climate change and habitat degradation (Wan *et al*. 2021) and are threatened by multiple causes, including deforestation, use of fuel-wood and charcoal, commercial logging, and collection and consumption of fruit and seeds (Harper et al., 2007). Time-series analysis of Landsat images empirically showed that Madagascar has lost 4.85 million hectares of tree cover since 2000, equivalent to a 25% decrease in tree cover (Suzzi-Simmons, 2023). These threats, along with low regeneration rates and the effects of climate change, pose a risk of ecological extinction for many plants in Madagascar like the baobabs trees (Wilson, 2018). The loss of tree cover due to deforestation contributes to the overall decrease in habitat for baobab, further endangering their population. Recent studies (Wan *et al*. 2021; Vielledent *et al*. 2013) predict the potential extinction of *Adansonia suarezensis* by 2080 due to shrinking suitable habitats and suggesting this species as critical endangered. The authors showed that there is a considerable population size reduction for the populations of this species. *A. suarezensis* is one of the species occurred in the north of Madagascar with restricted current distribution area around 1200km^2^ (Vieilledent *et al*. 2013). Efforts to protect and restore the forests, as well as sustainable management practices, are crucial for the conservation of baobabs and their habitat in Madagascar.

The species *A. suarezensis* belongs to the Brevitubae section. This species is a large, hermaphroditic tree that can grow up to 25 meters tall and 2 to 5 meters in diameter. Its shape is typically ‘T’ or ‘Y’ shaped with no secondary branching. The bark is smooth, gray to reddish-gray in color. The flowers are white with numerous white stamens. The fruit is an oblong pod, covered in a leathery exocarp called a pod, which is brown and pubescent. The fruit contains several kidney-shaped seeds, with a notable invagination (Baum, 1995a). *A. suarezensis* pollination was documented as bats’ pollination and some species of hawkmoth (Baum 1995 b; Ryckewaert *et al*. 2011).

Baobab genetic structure, population genetics and factors influencing genetic diversity have been studied in several papers. Leong Pock Tsy et al. (2013) and Karimi et al. (2022) highlighted genetic differentiation among baobab species, emphasizing their adaptation to diverse climates and habitats in Madagascar. However, the lack of data regarding gene flow, genetic drift, and inbreeding within and between *A. suarezensis* populations raises questions about the species’ evolutionary dynamics and adaptation potential in northern Madagascar making it hard to understand their evolutionary potential and adaptation strategies.

Wiehle *et al*. (2014) analyzed the genetic and morphological variability of East African populations of *Adansonia digitata* in Sudan. They found balanced genetic diversity that did not differ between locations or management regimes suggesting a positive effect of human intervention. Rangan et al. (2015) investigated the role of human agency in the gene flow and geographical distribution of the Australian baobab, *Adansonia gregorii* and found weak geographic structure and high gene flow, with humans identified as the most likely dispersal vectors. Chládová *et al*. (2019) found high genetic diversity and significant differences between coastal and inland populations in *A. digitata*, suggesting limited gene flow between them. *A. suarezensis* had lower genome wide heterozygosity and numerous segments of long runs of homozygosity suggested high levels of recent inbreeding (Wan *et al*. 2024).

Inbreeding within and between *A. suarezensis* populations raises important questions about the species’ evolutionary dynamics and adaptation potential in northern Madagascar, complicating our understanding of their evolutionary capabilities and adaptation strategies. To evaluate the high levels of inbreeding, we propose identifying data on genetic drift in this study.

Our study addresses the following research questions: What is the genetic structure within and between populations of *A. suarezensis*? Does genetic drift influence the genetic diversity of *A. suarezensis* populations? Is there evidence of inbreeding within *A. suarezensis* populations? What are the relationships between plant population size, fitness, and genetic variation in *A. suarezensis*?

Understanding the genetic diversity, structure, and adaptation of *A. suarezensis* is essential for developing effective conservation strategies to ensure its long-term survival in Madagascar. The insights from our study can also inform conservation efforts for other vulnerable baobab species.

## Material and methods

### Study site

The investigation was conducted in the northernmost part of Madagascar, within the Antsiranana province, encompassing the DIANA region spanning an area of 44,784 km^2^ (**Figure 1**). The latter area includes unique habitat that hosts *A. suarezensis* and is characterized by falls within lowland dry forest ecotone (Humbert, 1965). Climatically, this region experiences concentrated rainfall from November to April with an average annual rainfall of circa 792mm. The mean annual temperature in the region is between 25.4°C and 26°C, with the maximum monthly rainfall recorded at 247 mm in January.

**Figure 1.**
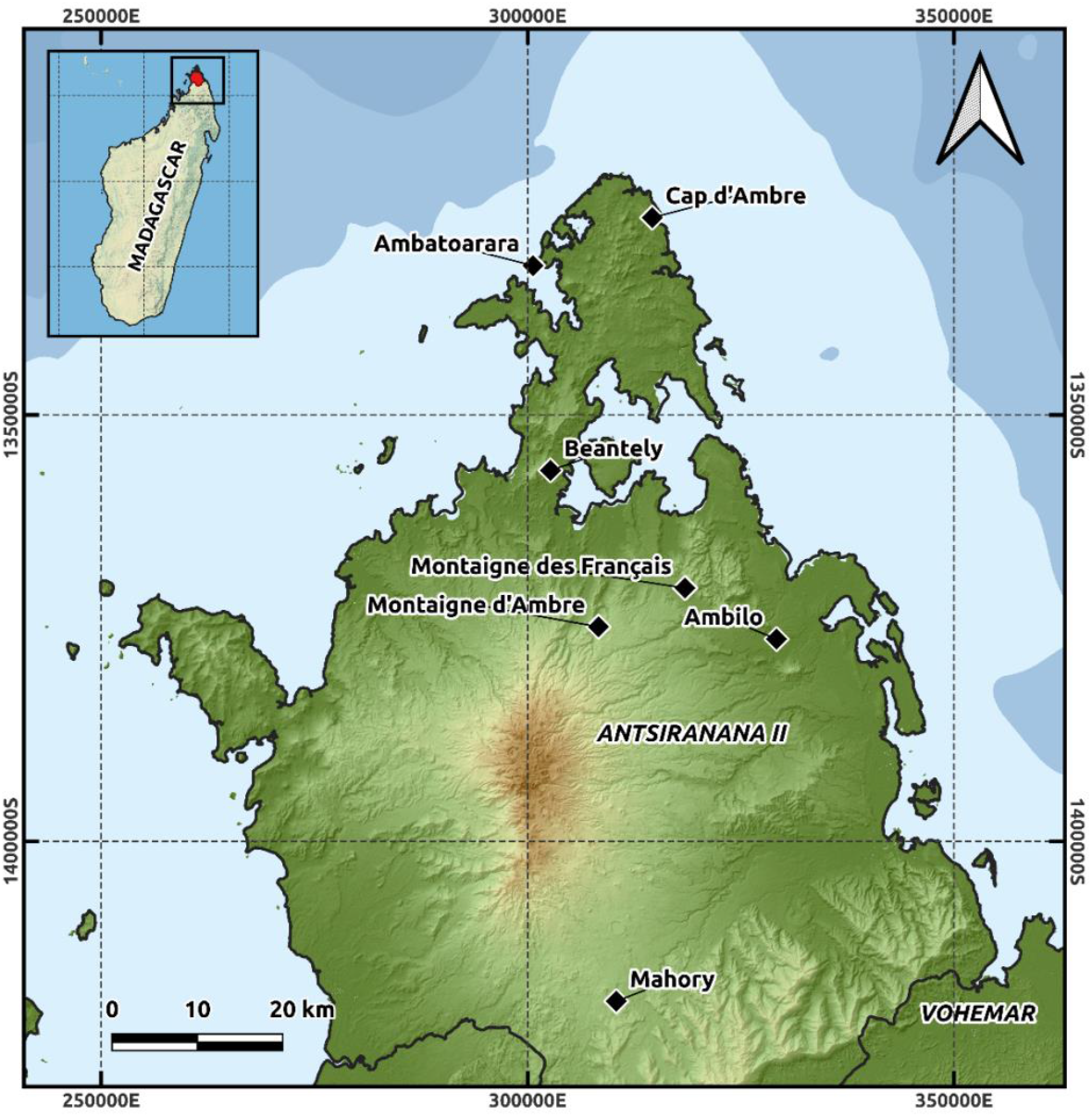
Geographic location of the seven analyzed populations.

Genetic data collection was carried out across seven populations. Two populations, Montagne d’Ambre and Ambatoarara, were newly identified during this study. This discovery addresses important gaps in the previously documented distribution of *A. suarezensis*, contributing to a more comprehensive understanding of its spatial patterns and potential conservation requirements.

A total of 419 individual trees were sampled across seven populations (Mahory, Ambatoarara, Cap d’Ambre, Montagne d’Ambre, Motaigne de Francais, Beantely, Ambilo) (**Table 1**). For each tree, one to two healthy leaves were collected. Samples were bagged, labeled, kept dry with silica gel for genetic analysis.

**Table 1.** Labels, geographic locations, and GPS coordinates of the seven *A. suarezensis* populations.

| Collection site | Label | N | Average GPS reading |  |
| --- | --- | --- | --- | --- |
|  |  |  | latitude_South | longitude_Est |
| Cap d'Ambre | CAPD | 25 | S 11°59'52.0" | E 049°17'49.5" |
| Ambilo | AMBL | 75 | S 12°26'42.4" | E 049°25'42.1" |
| Beantely | BEAT | 61 | S 09°42'01.2" | E 053°18'04.5" |
| Montaigne des Français | FRAN | 50 | S 12°23'25.5" | E 049°19'48.2" |
| Montaigne d'Ambre | MOTD | 25 | S 12°25'51.1" | E 049°14'11.0" |
| Ambatoarara | RARA | 25 | S 12°02'51.8" | E 049°10'06.5" |
| Mahory | MAHO | 158 | S 12°49'39.9" | E 049°15'10.5" |
| Total |  | 419 |  |  |

### DNA extraction and SSR analyses

Genomic DNA was extracted from young baobab leaves using a modified CTAB method. Fresh leaf tissue (approximately 200 mg) was subjected to mechanical disruption in a 2.0 mL microcentrifuge tube containing a single 5 mm zirconia bead using a mixer mill (Qiagen MM300) at 30 Hz for 60 seconds. To remove secondary metabolites, a washing step was performed three times prior to CTAB extraction. The washing buffer consisted of 0.1 M HEPES-HCl (pH 8.0), 1.02 % PVP (polyvinylpyrrolidone, MW 40,000) (w/v), 0.9 % L-ascorbic acid (w/v), and 2 % 2-mercaptoethanol (v/v). Subsequent DNA purification was performed using the DNeasy Plant Mini Kit (Qiagen) according to the manufacturer’s protocol.

### Genome-wide nuclear markers

To investigate the genetic diversity and population structure, we employed the MIG-seq method (Suyama and Matsuki, 2015), which targets single nucleotide polymorphic (SNP) sites and generates a comprehensive dataset suitable for population demographic analyses. Briefly, MIG-seq amplifies thousands of genome-wide regions using multiplexed inter-simple sequence repeat (ISSR) primers for PCR amplification. This approach bypasses the conventional steps of genomic DNA fragmentation and ligation of barcoded adapters, thereby reducing the cost and labor associated with library preparation. For this study, we utilized MIG-seq primer set-1 developed by Suyama and Matsuki (2015). The MIG-seq library was prepared following the protocol described by Suyama et al. (2022) and sequenced using the MiSeq system (Illumina, San Diego, CA, USA) with the MiSeq Reagent Kit v3 (150 cycles).

### Bioinformatic and population structure analyses

Out of the 161 samples analyzed, 8 samples were excluded due to an insufficient number of forward or reverse raw reads (<3000). The MIG-seq raw reads were trimmed using Trimmomatic v0.32 (Bolger et al., 2014) with the following parameters: HEADCROP:6, CROP:77, SLIDINGWINDOW:10:30, and MINLEN:51. After trimming, the reads were merged and processed together. Since the insert size of the sequenced molecules were longer than the number of cycles sequenced, no overlap between the forward and reverse reads was expected. Genotypes were called using the STACKS v2.6.1 pipeline (Rochette et al., 2019). The minimum depth for genotype calling was set to 6 (-m 6), while default values were applied for the other parameters. In the STACKS pipeline, cstacks was used to build a catalogue, followed by sstacks to align reads to the catalogue and generate stacks. The tsv2bam function was employed to transpose the genotype data by locus into BAM files, which were subsequently analyzed using gstacks in “De novo mode” to call genotypes. Finally, the populations module was used to extract SNPs with appropriate genotyping rates for each analysis, excluding loci with high observed heterozygosity (>0.6) and SNPs with a minor allele count of less than 3. To construct a maximum likelihood tree, RAxML (Stamatakis, 2014) was employed with the GTRGAMMA model using SNPs with a genotyping rate >10% (-R 0.1). Summary statistics were calculated using the populations program implemented in Stacks2. The analyses were performed using SNPs with a genotyping rate >80% (-R 0.8). To evaluate population structure, the ADMIXTURE program (v1.3.0) (Alexander et al., 2009) was used with k values ranging from 2 to 10. Cross-validation (CV) error was computed to identify the optimal value of k that provided the highest predictive accuracy. For the ADMIXTURE analysis, SNPs with a genotyping rate >80% were used after removing highly linked SNPs (R^2^ > 0.4) with PLINK v1.9 (Chang et al., 2015).

## Results

### Genetic structure between populations

The results of the analysis of the seven *A. suarezensis* populations revealed significant and distinct genetic structures. Admixture analysis based on 1374 SNPs (best K=7) and 533 SNPs (best K=9) consistently demonstrated two primary genetic clusters. The Mahory population (MAHO) formed a distinct genetic cluster, separate from the other six populations, regardless of the number of clusters modeled (e.g., K=2 to K=9; **Figure 2**).

**Figure 2.**
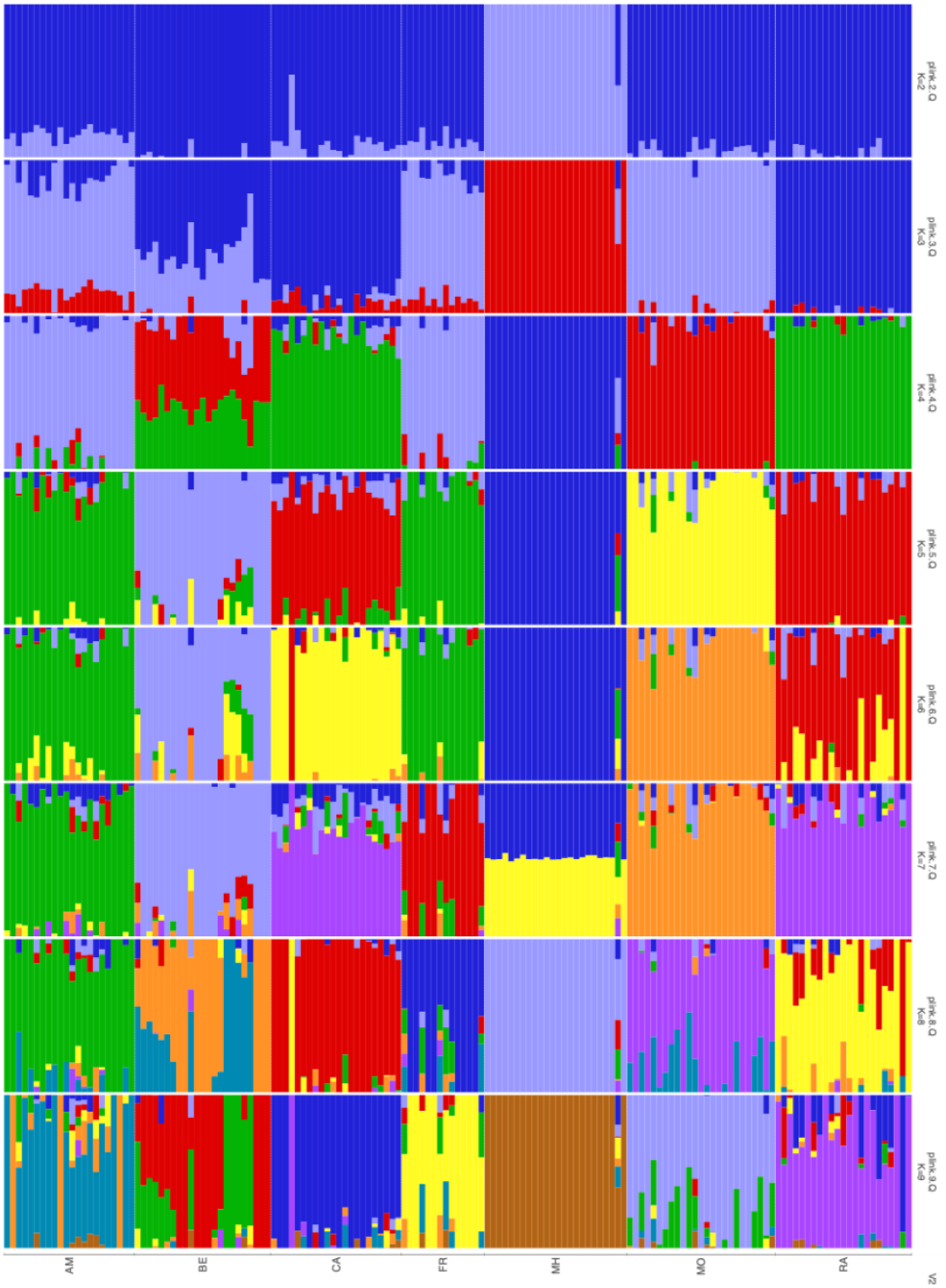
Admixture analysis from K=2 to K=9. Each vertical bar represents an individual, and colors indicate genetic ancestry proportions assigned to different clusters.

This genetic distinction suggests that Mahory population represents a unique and valuable genetic resource within *A. suarezensis*. In contrast, the other populations, including Cap d’Ambre (CAPD), Ambilo (AMBL), Beantely (BEAT), Montagne des Francais (FRAN), Montagne d’Ambre (MOTD), and Ambatoarara (RARA), exhibited varying degrees of genetic admixture, with evidence of some gene flow among northern populations. The phylogenetic tree constructed using 3856 SNPs demonstrated clear genetic relationships between populations. Mahory consistently appeared as a distinct clade, reinforcing its genetic isolation and limited gene flow with the other populations (**Figure 3**). Northern populations showed more interconnected relationships, with minor sub-clusters reflecting varying levels of shared genetic patterns. Only the two populations of Cap d’Ambre and Ambatoarara are very closely related, indeed there are two individuals that mixes together, probably sharing similar genetic makeup. Then, other groups of populations are composed by individuals from MOTD and BEAT populations, and individual from AMBL and FRAN populations.

**Figure 3.**
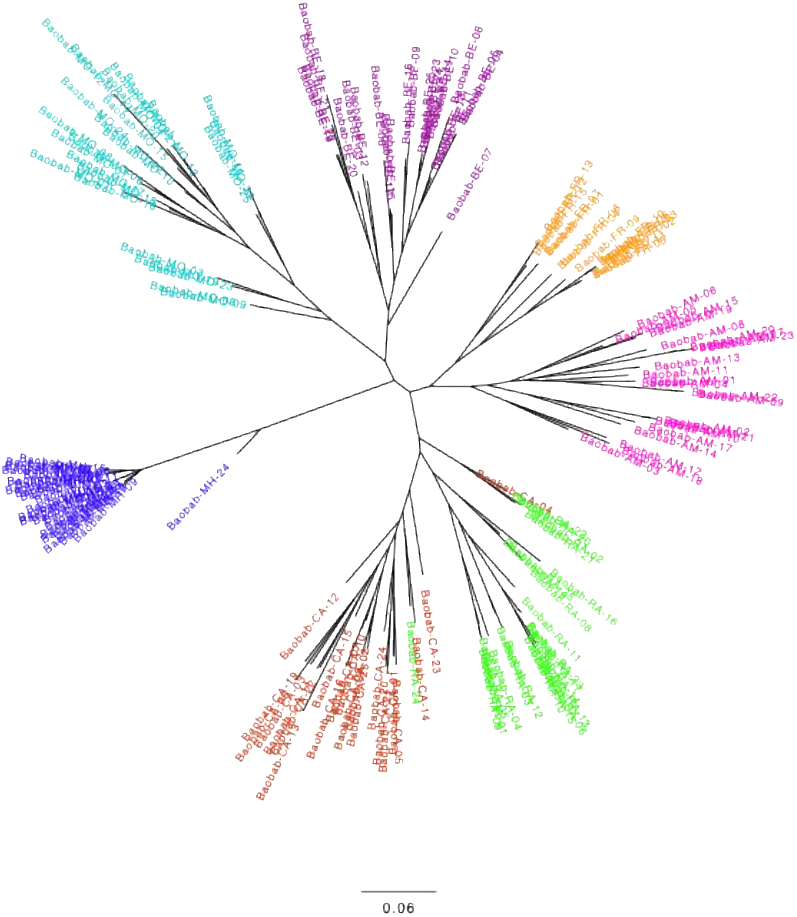
Phylogenetic tree based on 3,856 SNPs showing genetic relationships between populations. The tree is constructed using Maximum Likelihood, with branch lengths representing genetic distances.

### Phylogenetic Relationships

#### Genetic relationship among populations

The genetic diversity metrics derived from the 657 SNPs dataset indicated moderate levels of diversity across populations. Observed heterozygosity values ranged from 0.175 in Mahory to 0.212 in Montagne d’Ambre. Similarly, nucleotide diversity (π) ranged from 0.103 in Mahory to 0.224 in Ambilo. Inbreeding coefficients (Fis) varied between populations. Cape d’Ambre population has the highest value of inbreeding (0.079) whilst the Mahory population has a negative value of inbreeding coefficient (-0.112).

The analysis using the 1800 SNP dataset reinforced these patterns showing slighly higher resolution in the genetic diversity estimates. The nucleotide diversity (π) has values ranging from 0.11 of the Mahory population to 0.221 of the Ambilo population. Regarding the inbreeding coefficient there are more relevant differences. In fact, the two population Ambilo and Cape d’Ambre have the highest values of inbreeding, 0.146 and 0.125 respectively. On the other side, the Mahory population has higher values than before (-0.065).

An additional analysis was performed to compare diversity estimates based on variant positions alone versus all positions (variant and fixed). The inclusion of fixed positions resulted in a slight decrease of information captured from the invariant loci. Noticeably, we observed that from the 657 SNPs dataset the percentage of polymorphic sites in the Mahory population is much lower than the other (35%) and the same happened in the 1800SNP dataset where the percentage is 43%.

#### Admixture Patterns

The admixture analysis provides insights into the ancestral composition of *A. suarezensis* populations. Using 1374 SNPs (best K=7), the populations displayed distinct ancestral contributions. The Mahory population exhibited a unique ancestry pattern, consistent with its genetic isolation. Other populations, such as Beantely (BEAT) and Cap d’Ambre (CAPD), displayed significant admixture, indicating contributions from multiple ancestral populations.

The analysis based on 533 SNPs (best K=9) offered a finer resolution, revealing additional substructures. For instance, individuals within northern populations displayed more heterogeneous ancestry patterns, suggesting historical migration or gene flow events. The colored bars in the admixture plots illustrate the proportion of ancestral contributions for each individual. While populations such as Ambatoarara (RARA) and Montagne d’Ambre (MOTD) exhibited mixed ancestry patterns, Mahory remained distinct with minimal signs of admixture. This further underscores its unique genetic makeup.

## Discussion

The population genetics analysis of *A. suarezensis* showed a division into two distinct groups based on their geographical origin, similar to findings from studies on *A. digitata*. In the study of Chládová et al. (2019), conducted in southeastern Kenya, Bayesian clustering and Principal Coordinate Analysis (PCoA) indicated a separation between inland and coastal populations. This separation aligns with observations of distinct inland and coastal clusters in other baobab studies conducted in different regions, such as Benin and East and southeast African countries (Assogbadjo et al., 2006; Munthali et al., 2013). Several authors suggested that climate, human factors, pollination, seed dispersal and elevation differences might contribute to this population genetic differentiation between baobab populations. For *A. suarezensis*, a similar inland-coastal pattern is evident.

The Mahory population’s genetic distinctiveness may result from its distance from the other populations in the region (thus to be considered as an inland population) leading to a geographic isolation, with the Montagne d’Ambre massif acting as a biogeographical barrier and its location in a more humid environment compared to the drier northern populations. This isolation, coupled with a negative inbreeding coefficient, suggests an outbreeding strategy maintaining high heterozygosity. Such heterozygosity is likely to confer enhanced genetic resilience, enabling adaptability to environmental fluctuations and ecological pressures (Sellis et al., 2011; Hedrick et al., 2012). However, Mahory’s low proportion of polymorphic sites raises concerns about its adaptability to changing conditions. This apparent paradox likely reflects historical bottlenecks or founder effects that reduced allele diversity, balanced by effective outbreeding mechanisms. For instance, Munthali et al. (2013) found low genetic diversity in *A. digitata* populations in Malawi, with a percentage of polymorphic loci equal to 48%, attributed to marginal growing conditions, anthropogenic influences, and founder effects. Similar selective pressures may operate in Mahory, where natural selection in its unique environment could favor heterozygosity, conferring resilience to ecological changes.

Conversely, northern populations exhibit a more interconnected genetic structure, with ongoing gene flow maintaining genetic diversity and higher numbers of polymorphic sites. However, high inbreeding coefficients in coastal populations like Ambilo and Cap d’Ambre suggest localized inbreeding depression, potentially due to habitat fragmentation or reduced effective population sizes and the admixture analysis supports these patterns. Mahory exhibits minimal gene flow with northern populations, contrasting with the significant admixture observed among northern populations. This connectivity highlights the dynamic interplay of genetic drift and environmental adaptation shaping diversity. Additionally, pollinators and seed dispersers play a crucial role in facilitating gene flow and enhancing genetic connectivity among populations. Comparative pollination studies across populations could further elucidate these patterns.

Historical climatic changes and human interventions may also contribute to the observed genetic differentiation. Climatic shifts during the late Holocene (around 3700 years BP) likely influenced baobab distributions, favoring coastal adaptations and subsequent adaptation to more humid conditions (Chládová et al., 2019). Moreover, in the work of Wan et al. (2024), they modelled the distribution of different baobab species under different climate scenarios, and during the last glacial maximum (∼22.000 years before present) *A. suarezensis* had more extensive distribution in the northern part of Madagascar, indicating a more continuous range and not fragmented as it is now.

Alternatively, human-mediated introductions of baobabs to coastal regions may have established distinct ecotypes over time. Anthropogenic dispersal in northern regions could explain the observed admixture and genetic connectivity, while Mahory’s distinctiveness suggests limited human influence, retaining a more natural evolutionary trajectory. This could be explained by the possibility that human populations planted some baobabs in the northern region suggesting a history of anthropogenic dispersal.

The phylogenetic analysis highlights Mahory’s distinct clade, emphasizing its genetic isolation, while relationships among northern populations, the populations from Montagne d’Ambre, Beantely, Ambilo, and Montagne des Français populations group together indicating shared ancestry or historical connectivity. Also Cap d’Ambre and Ambatoarara populations are very closely related, showing also mixing individuals in the clades, revealing shared ancestry and potential gene flow. However, as stated before, this gene flow is not corroborated by inbreeding analyses, which indicate inbreeding issues in Cap d’Ambre and Ambilo.

Despite the clear genetic differentiation between inland and coastal baobab populations, the exact reasons behind this phenomenon remain unclear. Further research is needed to investigate whether these species were or not planted by humans. This could involve techniques such as dating the trees, expanding sampling efforts to identify new populations of *A. suarezensis*, and examining potential morphological changes associated with genetic differentiation. These studies would help determine whether distinct baobab ecotypes exist, each uniquely adapted to the specific climatic and environmental conditions of their respective locations.

The observed genetic differentiation and diversity patterns have significant conservation implications. Mahory’s unique genetic makeup requires targeted conservation efforts, prioritizing habitat preservation and mitigation of human impacts to safeguard its evolutionary potential. For northern populations, strategies should enhance genetic connectivity, addressing high inbreeding through habitat corridors and ecological networks, promoting resilience to environmental changes. These differences highlight the varying evolutionary pressures and reproductive strategies across populations.

Understanding the drivers of genetic isolation and adaptive divergence will be critical for developing conservation strategies that ensure the long-term survival of *A. suarezensis* and its genetic resources.

